# Mathematical Preconditions for Existence of the Stock-Recruitment Function

**DOI:** 10.1101/2023.10.18.562878

**Authors:** Ute Schaarschmidt, Anna S. J. Frank, Sam Subbey

## Abstract

For marine species, several life stages link parents to their progeny (recruits), through a process referred to as recruitment. Current stock recruitment (SR) functions encapsulate this multi-stage relationship in a closed-form mathematical expression, which explicitly relates the biomass of parents, to the number of recruits. This functional relationship (between parents and recruits) is required for management purposes. No study, however, has investigated the conditions that validate the existence of the SR functions when all life history stages are incorporated into a model.

In this study, we represent the processes leading to recruitment by a stage-and age-structured discrete-time population dynamic model. We show that in general, a SR function does not correctly represent the parent-progeny relationship. A valid relationship must incorporate information across the complete life cycle over several time periods. For populations with simple life cycle history, a functional relationship may result, which is not necessarily consistent with current SR functions.

We present a brief discussion of the relevance of our results to effective management of fisheries.

## 1 Introduction

The life history of marine populations often comprises a series of distinct stages or stanzas (Paulik 1973), each characterized by a unique set of factors influencing survival (Nash 1998) (see Fig. 1).

**Fig. 1.**
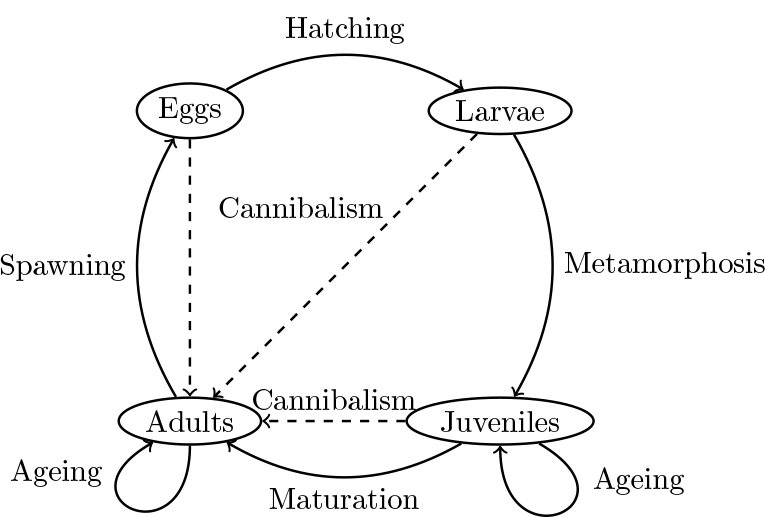
A schematic diagram of the life-history cycle of a fish with four stanzas. The principal developmental processes are indicated. Transitions from one age class to the next age class are represented by loops. In addition, the linkages between the adults and the early life-history stages through cannibalism are illustrated

These life stanzas encompass critical developmental phases such as eggs, larvae, juveniles, and adults. Principal developmental processes (e.g., spawning, hatching, metamorphosis, and maturation) cause transitions from one life stanza to the next (Paulik 1973).

Understanding this intricate dynamics of marine populations is fundamental for effective and sustainable fisheries management (to maintain healthy fish populations). Central to this understanding is the concept of stock recruitment (SR) and the SR function.

Fish stock recruitment refers to the relationship between the number of juvenile fish (recruits) that enter a population and the number of adult fish (the stock) that spawn to produce those recruits. It is a critical concept in fisheries biology and management because it helps (scientists and managers) understand and predict how changes in the number of adult fish (e.g., due to fishing or environmental factors) can affect the abundance of young fish that join the population. The stock recruitment relationship is often described by a function of one variable and defined in terms of numbers of individuals (Chambers and Trippel 1997). In fisheries science, the functions are usually unimodal and often assume that as the size of the parental population increases, the number of recruits also increases or remains constant (Beverton and Holt 1957; Ricker 1954). While these functions can be useful simplifications and can capture certain patterns, issues have been raised, which revolve around both their fundamental existence and formulation (Subbey et al 2014).

The SR function is conventionally assumed to exist. However, several studies have illuminated nuances in recruitment patterns, some of which are inconsistent with conventional SR functions (reviewed by Haddon (2011) and Hilborn and Walters (1992, chapter 7)). For instance, empirical multi-modal patterns of recruitment have been reported in the literature (Hennemuth et al 1980), which may only be explained by multi-modal fish SR functions. Such functions may be derived by considering multiple life stages prior to recruitment (Brooks and Powers 2007; Paulik 1973), though this is a departure from current convention. On the other hand, multi-stage model simulations have revealed that the existence of a stock-recruitment (SR) function may not be guaranteed under certain conditions. For instance, when parameters such as fecundity, reproductive rates, or predation rates vary with age, the traditional SR function may fail to emerge (Touzeau and Gouzé 1998). In contrast, in multi-stage models that account for recruitment as a function of an age-structured parental population, the resulting function becomes multivariate, diverging from the conventional SR function, which typically relies on a single variable (Schaarschmidt et al 2018). Thus, the quest of the paper is to determine necessary and sufficient conditions for existence of a (I) closed-form SR function, and (II) multivariate SR function.

In addressing the above issues, the paper uses theorems, which are summarized in a Ph.D thesis (Schaarschmidt 2018). This paper extends the work in Schaarschmidt (2018) by providing rigorous proofs and biological interpretations of the theorems. Furthermore, we discuss the choice of the mathematical model and interpretations of our results.

We will employ a systematic, modeling-driven approach to comprehend the survival and reproduction (SR) function within multi-stage population dynamics. In contrast to studies that rely on inherently variable and uncertain data to investigate SR relationships (Myers and Barrowman 1996; Gilbert 1997), our methodology consistently provides a mathematical representation of the parent-progeny relationship. Our approach involving research on multi-stage SR relationships is aligned with literature (Quinn and Deriso 1999; Schaarschmidt et al 2018; Touzeau and Gouzé 1998), but differ in the following way. In contrast to (Schaarschmidt et al 2018; Touzeau and Gouzé 1998), we consider all life history stages (eggs, larvae, juveniles and adults), rather than a broad classification of life stages into pre-recruits and recruits. We consider a more comprehensive life cycle process than in Quinn and Deriso (1999).

The article is organized in the following way. Section 2 presents the modeling framework adopted in this manuscript. It states the necessary assumptions and the equations of the general discrete time multi-stage model that is used to simulate the entire life history cycle of fish populations. Section 3 focuses on the parent-progeny relationship and its mathematical simplification into the multivariate Stock-Recruitment (SR) and closed-form SR-functions, by proving necessary and sufficient conditions for their existence. We also provide mathematical proofs of these conditions. In the discussion, Section 4, we compare our findings to current literature and provide biological interpretations for the necessary and sufficient conditions.

## 2 Modeling framework

An adopted modeling framework must satisfy certain criteria, which include the ability to clearly delineate different life stages, e.g., in Fig. 1. This specificity enables a more accurate representation of the dynamics at, and between each stage. While it is possible to categorize the adult population by length (Callahan et al 2019) or size (weight) (Meng et al 2013), we align with previous research on multi-stage SR relationships (Schaarschmidt et al 2018; Touzeau and Gouzé 1998), and focus on an age-structured adult population model.

We now specify a multi-stage model to describe the life-history cycle illustrated in Fig. 1, which is subject to the following biological (**B1–B2**) assumptions and model-time characteristics (**T1–T4**):

**B1**: Egg production varies with fecundity and the proportion of spawners.

**B2**: Mortality rates of larvae and juveniles are functions of numbers of adults, larvae, and juveniles.

The assumption **B1** follows a standard approach in fisheries (e.g. Hilborn and Walters 1992, chapter 3), while B2 reflects that survival may be affected by processes such as cannibalism, food availability, and competition (e.g. Hilborn and Walters 1992, chapter 7).

**T1**: Discrete Time Transitions: We assume discrete transitions between life stanzas (see e.g., Fig. 1).

**T2**: Unified Time Steps: We consider uniform time steps across all life stanzas.

**T3**: Time Delay Constraint: Our approach incorporates a positive time delay, recognizing the delay between spawning and recruitment.

**T4**: Transition Time: Spawning and transition to the juvenile stage are assumed to happen within one time step. If surviving, juveniles and adults age by one in every simulation time step.

The assumptions **T1, T2** and **T3** are consistent with the fisheries literature (Quinn and Deriso 1999, chapter 5). Assumptions **T1** and **T2** also allow for direct comparison between model predictions and observed data, as empirical data on marine populations often come in discrete time intervals. Assumption **T4** can be justified from the fisheries literature, with egg and larva stage duration of a few months, maturation after a few years, and life spans of several years (Petitgas et al 2013). Furthermore, faster evolution of prerecruits in comparison to adults is an assumption underlying other multi-stage models for SR (Schaarschmidt et al 2018; Touzeau and Gouzé 1998).

### A general discrete time multi-stage model (DTMM)

Based on **B1–B2** and **T1–T4**, we formally define an age-structured population dynamic model that describes births, survivals, and transitions from one stanza to the next. The considered stages are eggs (*e*), larvae (*l*), juveniles (*j*), and adults (*a*). We have age classes 0, …, *m*−1 representing juveniles and age classes *m*, …, *n* for adults. We use the symbols in brackets and 0, …, *n* as indices. An overview of the nomenclature for the model is given in Table 1.

**Table 1.** Nomenclature for the DTMM (adapted from Schaarschmidt (2018)).

|  |  |
| --- | --- |
| $\mathcal{S} = \{e, l, j, a\}$ | Set of life stages: eggs ($e$), larvae ($l$), juveniles ($j$), adults ($a$) |
| $m \in \mathbb{N}_0 = \{0, 1, 2, 3, \dots\}$ | Number of juvenile age classes |
| $n \in \mathbb{N}_0$ | Total number of age classes |
| $k \in \{0, \dots, m, \dots, n\}$ | Indices of age classes |
| $t \in \mathbb{N}_0$ | Simulation time |
| $\tau \in \mathbb{N} = \{1, 2, 3, \dots\}$ | Time delay |
| $E_t \in \mathbb{R}_{\geq 0}$ | Number of eggs at time $t$ |
| $L_t \in \mathbb{R}_{\geq 0}$ | Number of larvae at time $t$ |
| $J_{k,t} \in \mathbb{R}_{\geq 0}$ | Number of juveniles in class $k = 0, \dots, m-1$ at time $t$ |
| $\mathbf{J}_t = (J_{0,t}, \dots, J_{m-1,t})$ | Vektor with numbers of juveniles |
| $N_{k,t} \in \mathbb{R}_{\geq 0}$ | Number of adults in class $k = m, \dots, n$ at time $t$ |
| $\mathbf{N}_t = (N_{m,t}, \dots, N_{n,t})$ | Vektor with numbers of adults. |
| $R_t = N_{m,t}$ | Number of individuals included in stage $m$ at time $t$ |
| $p_k \in (0, 1)$ | Proportion of adult female population in class $k = m, \dots, n$ |
| $f_k \geq 0$ | Fecundity of adult fish in class $k = m, \dots, n$ |
| $s_i$ | Function representing probability of surviving stage $i \in \{e, l, 0, \dots, n\}$ in a time step |
| $e : \mathbb{R}_{\geq 0}^n \rightarrow \mathbb{R}_{\geq 0}$ | Function representing egg production in a time step |

Let 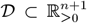, where *𝒟* is non-empty. The general discrete time multi-stage model (DTMM) is defined by (1)–(7) with initial condition (**J**_0_, **N**_0_) ∈ *𝒟*.

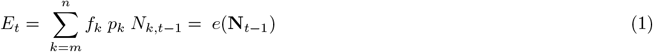

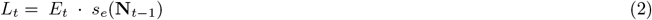

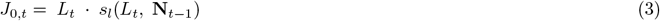

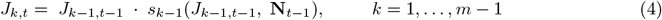

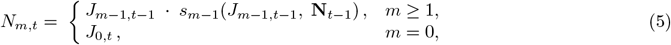

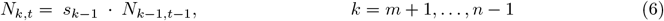

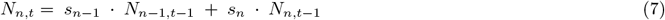

Most age- and/length structured marine populations have matured individuals distributed over several age- and length groups (see e.g., Jokar et al (2021)). Hence, total egg production in (1) is defined as the sum of eggs produced over all age classes and depends on the adult population. The number of larvae presented in (2) is the number of eggs that survive to the next time step. The juveniles may exist in several age categories (see (3) and (4)) and Fig. 1. The initial juvenile population of age 0 transitions directly from the larvae stage, and its survival may depend on the number of larvae and the adult fish population. The older juvenile populations of age *k* = 1, …, *m*−1 transition from the previous age group into the next, and their survival may be influenced by the number of juveniles within their own age group and the adult population. The rationale behind the survival rate is that (i) juveniles within each age group *k* compete for the same food resource and habitat (ii) the adult population could cannibalize on the lower developmental stanzas, i.e., larvae and juveniles. Maturation to the youngest age class of adults as described by (5) is either directly from the initial juvenile stage (if *m* = 0), or a juvenile group of age *m*−1 (if *m*≥1). As described by (6), the adults age linearly. The oldest age class of adults presented in (7) may consist of several age classes, i.e. individuals of any age≥*n*. All adult age classes *m*, …, *n* can produce eggs (see (1)).

The following conditions guarantee non-negative solutions of (1)–(7) with initial conditions 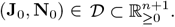 .

**Condition 1**. *For all k* = *m*, …, *n, let p*_*k*_ ∈ (0, 1) *and f*_*k*_ ≥ 0 *(as in Table 1) and*

**Condition 2**. *the codomains of functions s*_*i*_ *are subsets of* (0, 1].

Furthermore, we assume that 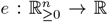 defined by (1) gives egg production as a non-constant function of **N**_*t*−1_, i.e., 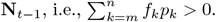.

## 3 The SR relationship in multi-stage population dynamics

Given the general DTMM in (1)–(7), we focus in this section on the parent-progeny relationship, as well as mathematical formulations describing this relationship in form of SR functions.

Recruitment *R*_*t*_ = *N*_*m,t*_ is defined as the number of individuals included in stage *m* at time *t* ∈ ℕ_0_, and represented in (5) in the general DTMM. Stock *S*_*t*_ is defined as an aggregation of the counts of adults represented by a weighted sum (see (10)).

While the parent-progeny relationship is the general term that describes the relationship between an adult population and their offspring, the traditional SR functions have been defined to mathematically describe the parent-progeny relationship. Whether the existence of SR functions is mathematically valid in a multi-stage framework has not been proven and is the goal of this section.

To establish conditions leading to a closed-form SR-function, we proceed in the following way:

**Step 1:** We use the DTMM (in (1)-(7)), representing the full life history cycle, to define the general parent-progeny relationship in Section 3.1.

**Step 2:** We then define and prove conditions under which the general parent progeny relationship can be simplified into a multi-variate SR-function (not yet closed-form) (see Section 3.2).

**Step 3:** In Section 3.3, we identify further conditions that allow us to reduce the multi-variate SR-function into a closed-from SR-function.

In order to establish conditions for step 2 and 3, we posit two hypotheses for the existence of SR functions, based on our general DTMM.

### Hypothesis 1.

∃*τ* ∈ [1, *m* + 1], 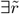 *such that for all t* ≥ *τ*,

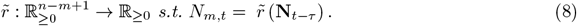

### Hypothesis 2.

∃*τ* ∈ [1, *m* + 1], ∃*r such that for all t* ≥ *τ*,

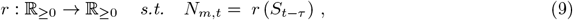

*where*

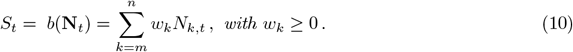

The functions 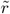 and *r* are the multivariate SR, and SR functions, respectively.

### Remark 1.

*Examples of S*_*t*_ *are the total number of adults (with w*_*k*_ = 1 *for all k* = *m*, …, *n) and the spawning stock biomass (the total weight of fish matured enough to contribute to the reproduction process). In the latter case, w*_*k*_ *>* 0 *is the average biomass of an individual in class k, respectively*.

### Remark 2.

*The requirement that τ* ∈ [1, *m* + 1] *stems from the observation that R*_*t*_ = *N*_*m,t*_ *is in general a function of* **N**_*t*−*m*−1_, …, **N**_*t*−1_, *as we will see in the following subsection*.

If the SR function *r* exists, we may define function 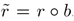, the composite function of *r* with the linear function *b* : ℝ_≥0_→ℝ_≥0_ defined by (10). Function 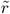 is a multivariate SR function.

### Remark 3.

*Existence of a multivariate SR function is a necessary condition for the existence of a SR function*.

### 3.1 General form of the parent-progeny relationship

The following lemma describes the fundamental link between adults and recruitment.

#### Lemma 1.

*Let E*_*t*_, *L*_*t*_, **J**_*t*_, **N**_*t*_ *for t* ≥ 0 *satisfy (1)–(7) with* (**J**_0_, **N**_0_) ∈ *𝒟. Then, R*_*t*_ = *N*_*m,t*_ *is given by (11) for all t*≥*m* + 1. *Here, the probability j*_*k*_ *of survival of eggs to class k* = 0, …, *m is defined recursively by (12)–(13)*.

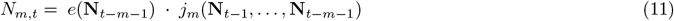

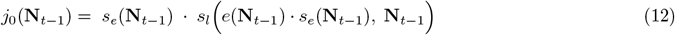

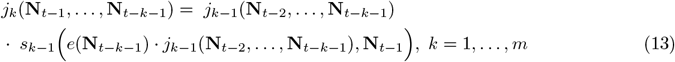

*Proof*. By (1)–(2), we can substitute *e*(**N**_*t*−1_)·*s*_*ϵ*_(**N**_*t*−1_) for *L*_*t*_ into (3). We then find *J*_0,*t*_ as a function of **N**_*t*−1_ given by (14). Using (12), we then substitute *j*_0_(**N**_*t*−1_) into (14) to find that (15) holds for *k* = 0.

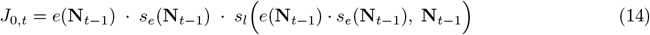

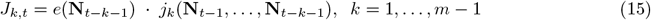

Assuming (15) to hold for *k* − 1 ∈ *{*0, …, *m* − 1*}*, we find

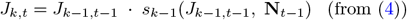

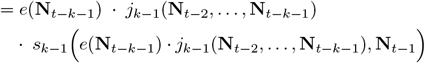

and thus, by definition (13), we obtain (15) for *k*. By induction, (15) holds for all *k* = 0, …, *m* − 1.

Analogously to one induction step, we use (5), (13) and (15) with *k* = *m*−1 to find *N*_*m,t*_ as given by (11). □

We thus observe that the DTMM describes recruitment *R*_*t*_ = *N*_*m,t*_ as a function of the vectors **N**_*t*−*m*−1_, …, **N**_*t*−1_ of numbers of adults at times *t*−*m*−1, …, *m*−1. In general, one needs to track the life cycle from the time of spawning to the time of recruitment and use information about each year’s adult population to obtain knowledge about the SR relationship. The general parent-progeny relationship involves a series of past states of the adult population at several time steps.

The general form of the parent-progeny relationship admitted by the DTMM is illustrated in Fig. 2. Egg production and the number of larvae and juveniles of age 0 in time step *t* − *m* are functions of **N**_*t*−*m*−1_. Numbers of juveniles of ages *k* ≥ 1 are given by functions of the numbers of juveniles in the previous year and age class and the numbers of adults in the previous time step. Recruitment *N*_*m,t*_ at time *t* is derived from the variables **N**_*t*−*m*−1_, …, **N**_*t*−1_.

**Fig. 2.**
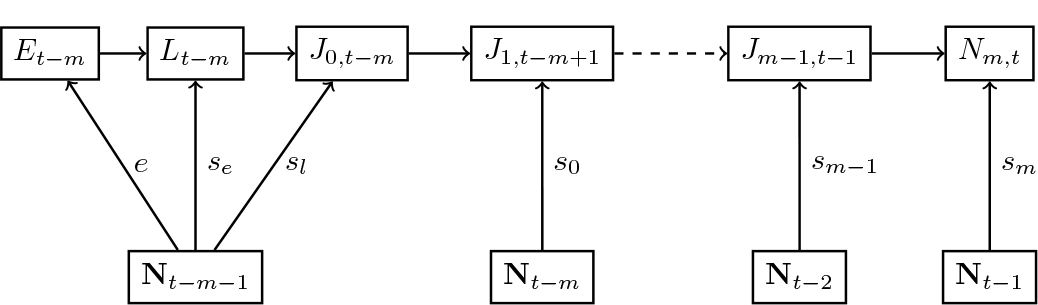
The general parent-progeny relationship for the DTMM. The evolution of individuals spawned at time *t*−*m* to recruitment at time *t* is given by (1)–(5). Here, the functional relationships given by the DTMM are represented by arrows. As an example, an arrow from *N*_*t*−*m*−1_ to *E*_*t*−*m*_ represents the assumption that egg production at time step *t* − *m* is given by function *e* of **N**_*t*−*m*−1_ (From Schaarschmidt (2018))

Using the recursive definition for the probability of survival of eggs to recruitment, we may redefine the multi-stage model in terms of fewer variables.

#### Corollary 1.

*Let E*_*t*_, *L*_*t*_, **J**_*t*_, **N**_*t*_ *for t* ≥ 0 *be a solution of a DTMM. Then*, **N**_*t*_ *with t* ≥ 0 *is a solution of a dynamical system given by (11)–(13) and (6)–(7) with initial condition* 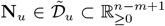, *for suitable* 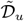 *and u* = 0, …, *m*.

The alternative formulation of the DTMM in Corollary 1 involves only one stage (adults), but the difference equation is of order *m*. We use Corollary 1 to prove the following result.

#### Lemma 2.

*Let E*_*t*_, *L*_*t*_, **J**_*t*_, **N**_*t*_ *for t* ≥ 0 *be a solution of a DTMM and* **N**_*t*_ = **N** *for all t* ∈ [0, *m* + 1]. *Then*, **N**_*t*_ = **N** *for all t* ≥ 0.

*Proof*. From (11) and **N**_*t*_ = **N** for *t* = 0, …, *m* + 1, it follows that *N*_*m,m*+2_ = *N*_*m,m*+1_. With (6)–(7) and **N**_*m*+1_ = **N**_*m*_, we obtain *N*_*k,m*+2_ = *N*_*k,m*+1_, for all *k* = *m* + 1, …, *n*. Overall, we found that **N**_*t*_ = **N** for *t* = 0, …, *m* + 1 implies **N**_*t*_ = **N** for *t* = 0, …, *m* + 2 and thus, by induction, **N**_*t*_ = **N** for all *t* ≥ 0.

In other words, having a constant adult population for a duration that corresponds to the delay between spawning and recruitment implies the population to be constant for all future times.

### 3.2 Existence of a multivariate SR function

In this subsection, we investigate, whether the multi-stage model always admits a multi-variate SR function, and state sufficient conditions for the existence of the multi-variate function.

#### Theorem 1.

*There exist DTMMs without multivariate SR function (as defined in Hypothesis 1)*.

The proof is by construction, using the following example of a DTMM.

#### Example 1.

*We consider a dynamical system of form (1)–(7) under assumptions (16)–(19). Let 𝒟 be a cuboid with positive volume, m* = 1 *and n* = 2.

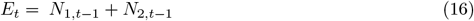

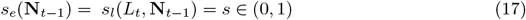

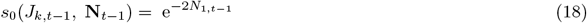

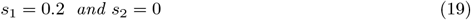

*Then, the dynamical system is given by (20)–(22) with (21) for R*_*t*_ = *N*_1,*t*_. *It is an example of a DTMM, since conditions 1–2 hold*.

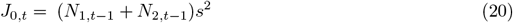

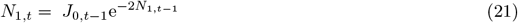

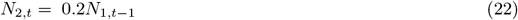

#### Proof of Theorem 1.

Our goal is to show that there exist solutions to the dynamical system that are such that a multivariate SR function cannot exist for any *τ* ∈ [1, *m* + 1] = [1, 2].

Let 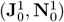, 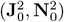 ∈ 𝒟 such that 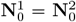 and 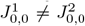. These vectors exist in *𝒟* since the cuboid is assumed to have positive volume. Denote by 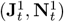 and 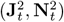 the solutions to (20)–(22) with initial condition 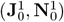 and 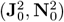, respectively.

**The case *τ* = 1**. For this case, we let *t* = 1 and consider recruitment *N*_1,1_ as given by (21). With the assumptions 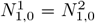 and 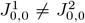 about the initial conditions, we see that 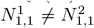 . As 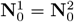 but 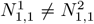, a multivariate SR function 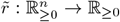 s.t. 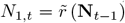 for all *t* ≥ 1 cannot exist.

**The case *τ* = 2**. Now, let *t* = 2. We substitute *J*_0,0_ as defined by (20) into (21) for *N*_1,*t*_ and obtain (23) for recruitment at time *t* = 2. Since 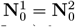 and 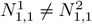, we observe that 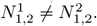 . Thus, a function 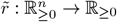 s.t. 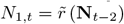 for all *t* ≥ 2 cannot exist.

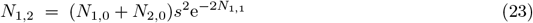

Summarizing, we have now shown that for *τ* = 1 and *τ* = 2, the DTMM (20)–(22) has solutions, for which there cannot exist any function 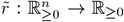 s.t. 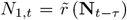 for all *t* ≥ *τ* . Hypothesis 1 does not hold for Example 1.

In the following, we consider a set of properties of Example 1. We consider them as logical statements and use terminology described e.g. in Rosen (2012).

#### Propositions

Define the following set of propositions, which are said to be true if they are true for all solutions

*E*_*t*_, *L*_*t*_, **J**_*t*_, **N**_*t*_, *t* ≥ 0 of a DTMM.

(C1) **N**_*t*_ = **N**_0_ for all *t* ∈ [0, *m* + 1],

(C2) *s*_*k*_(*J*_*k,t*_, **N**_*t*_) = *s*_*k*_(*J*_*k,t*_) for all *k* = 0, … *m* − 1 and for all *t* ≥ 0,

(C3) *m* = 0, i.e. *N*_*m,t*_ = *J*_0,*t*_ is given by (3).

A biological interpretation of (C3) is that there is only one cohort of juveniles. Then, recruitment of fish spawned at time *t* occurs at time *t* + 1.

**Remark 4**. *For Example 1, the logical solution of ((C1)* ∨ *(C2)* ∨ *(C3)) is false*.

*Proof*. The verification that (C1) evaluates as false follows directly from the proof of Theorem 1. Since there exists 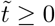 such that 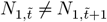 and (18) holds, (C2) cannot hold. The negation of (C3) holds by definition of the dynamical system. □

In other words, the multi-stage model from Example 1 without multivariate SR function has none of the properties (C1), (C2) or (C3). In the following, we will show that all three propositions individually ensure the existence of a multi-variate SR function.

#### Sufficient conditions for the existence of a multivariate SR function

##### Theorem 2.

*For all DTMMs, the logic solution of ((C1)* ∨ *(C2)* ∨ *(C3)) is sufficient for the existence of a multivariate SR function (Hypothesis 1). Function* 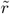 *is given by (24) and τ* = *m* + 1.

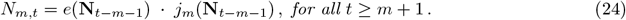

Here, we consider Lemma 1 and (11)–(13) given therein. For the convenience of the reader, the equations are re-stated here,

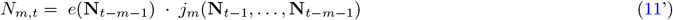

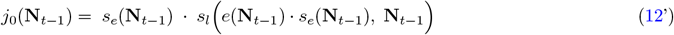

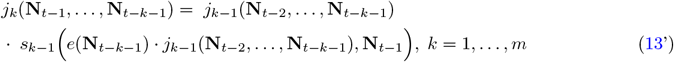

*Proof*. Assume that (C1) holds. With Lemma 2, we obtain that **N**_*t*_ = **N** for all *t* ≥ 0. By (11) for recruitment in Lemma 1, we can conclude that *N*_*m,t*_ is given by (24), for all *t* ≥ *m* + 1.

Using the recursive definition (12)-(13) for probability *j*_*k*_(**N**_*t*_), we observe that (C2) implies by induction that *j*_*k*_ is given by (25) for *k* = 1, …, *m* and *t* ≥ *k* + 1. In this case and due to (11), we obtain (24), for all *t* ≥ *m* + 1.

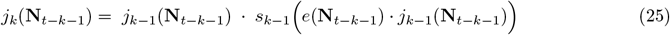

Proposition (C3) and *m* = 0 implies the existence of a multivariate SR function of form (24) for all *t* ≥ *m* + 1 by (11).

The simplification of the general parent-progeny relationship from Fig. 2 into a multi-variate SR function is illustrated in Fig. 3. The biological interpretation of Theorem 2 is that we can be sure to have a multi-variate SR function, if either

- the system is at a stable fixed point,
- survival rates of juveniles are constant with respect to the numbers of adults, or
- there is no more than one time step delay between spawning and recruitment.

**Fig. 3.**
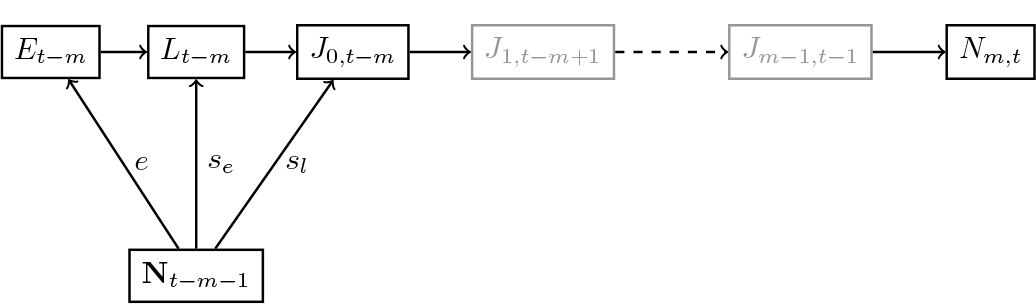
The multi-variate SR function for the DTMM. Here, the evolution of individuals spawned at time *t* − *m* to recruitment at time *t* is given by equations (1)–(5) under condition (C2)∨(C3). The resulting parent-progeny relationship is a multi-variate SR function (as recruitment is a function of **N**_*t*−*m*−1_). The gray color of some nodes represents the fact that under (C3), i.e., *m* = 0, the nodes *J*_1,*t*−*m*+1_ and *J*_*m*−1,*t*−1_ are not included in the graph (adapted from Schaarschmidt (2018)).

### 3.3 Existence of a SR function

We now state sufficient conditions for a multi-variate SR function to have the characteristic form of a SR function and investigate, what happens, if any of the conditions are violated.

For the DTMM, we define the following propositions. Here, we use linear function *b* : ℝ_≥0_ →

ℝ_≥0_ defined by (10).

(C4)(a) 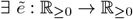 s.t. 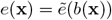, for all 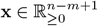,

(C4)(b) 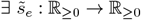 s.t. 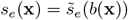, for all 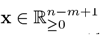,

(C4)(c) 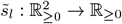 s.t. 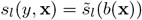 for all 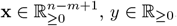, *y* ∈ ℝ_≥0_.

**Remark 5**. *Let E*_*t*_, *L*_*t*_, **J**_*t*_, **N**_*t*_, *t* ≥ 0, *be a solution of a DTMM and assume that (C1) is true. From Lemma 2, we know that (C1) implies N*_*m,t*_ = *N*_*m*,0_ *for all t*≥0 *and Hypothesis 2 is true*.

#### Theorem 3.

*Consider a DTMM, for which the logic solution of ((C2)* ∨*(C3)) is true. Then, violation of any of the logical statements (C4)(a)–(c) can imply that no SR function exists*.

*Proof*. Consider a DTMM such that (C4)(b) and (C4)(c) hold and (C4)(a) is violated.

Assume *m* = 0, i.e. a multivariate SR function exists. From the proof to Lemma 1, we know that recruitment *N*_*m,t*_ is given by (14), for *t*≥1. With (C4)(b) and (C4)(c), assuming the existence of a SR function would imply (26)–(27) for all *t*≥1. This would be a contraction to (C4)(a) being false. Thus, this DTMM cannot have any SR function.

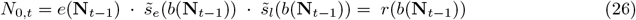

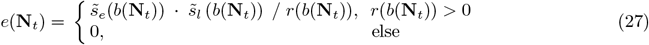

Now we consider the case *m >* 0 and assume the logical solution of (C2) is true. From (C4)(b) and (C4)(c) and (14), we see that *J*_0,*t*+1_ is given by (28), for *t* ≥ *m* + 1. Due to (C2) and (4)–(5), we obtain (29)–(30) and see that *N*_*m,t*_ is given by a function of *J*_0,*t*−*m*_. Since we also have *N*_*m,t*_ = *r*(*b*(**N**_*t*−*m*−1_)), we see that *J*_0,*t*_ must be given by a function 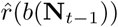. Then, we obtain (31) and a contradiction to the assumption that (C4)(a) evaluates as false.

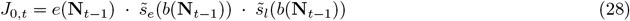

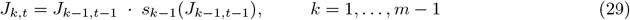

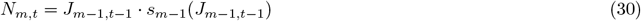

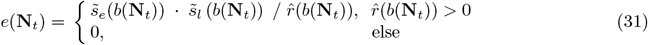

Analogously, we can define DTMMs, for which either (C4)(b) or (C4)(c) is violated, but (C4)(a) holds and there exists no SR function.

We have now proved that a multi-variate SR function is not required to have the characteristic form of a SR function unless we impose structural conditions on the functions representing egg production and survival rates of eggs and larvae. However, we will now see that the combination of these three conditions ensures the existence of a SR function. □

#### Sufficient conditions for the existence of a SR function

##### Theorem 4.

*For all DTMMs, the logic solution of ((C4)*∧ *((C2)* ∨ *(C3)))* ∨ *(C1) is sufficient for the existence of a SR function r given by (32) with τ* = *m* + 1.

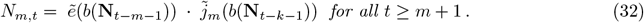

*Proof*. Let (C4) ∧ ((C2) ∨ (C3)) be true and *E*_*t*_, *L*_*t*_, **J**_*t*_, **N**_*t*_, *t* ≥ 0, a solution of a DTMM. Then, by Theorem 2 and Lemma 1, *N*_*m,t*_ is given by (12) and (24)–(25), for *t* ≥ *m* + 1. With (C4), we get *j*_0_(**x**) as a function of *b*(**x**) and given by (33), for all 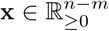. By induction and (25), we obtain (34) for all **x** ∈ ℝ ^*n*−*m*^ and *k* = 1, …, *m*. From Theorem 2, we see that the SR function is in this case defined by (32).

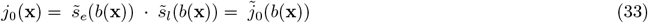

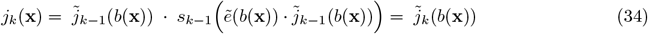

The existence of a SR function, if (C1) is true, follows directly from Remark 5.

The simplification of the general parent-progeny relationship for the DTMM into a closed-form SR function is illustrated in Fig. 4.

**Fig. 4.**
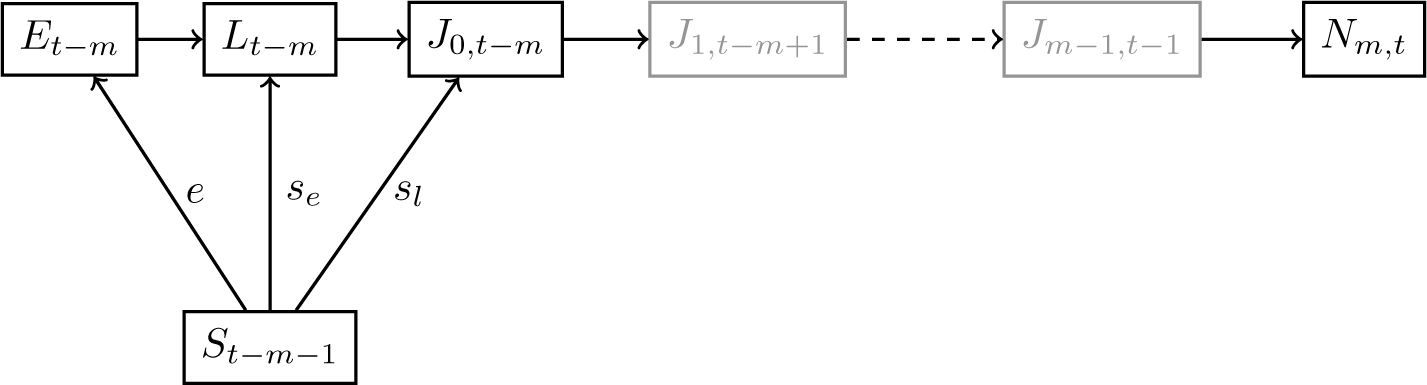
The closed-form SR function for the DTMM. The evolution of individuals spawned at time *t* − *m* to recruitment at time *t* given is by equations (1)–(5) assuming (C4) ∧ ((C2) ∨ (C3)) to be true. The resulting parent-progeny relationship is a closed-form SR function (since *N*_*m,t*_ is a function of *S*_*t*−*m*−1_). The gray color of some nodes represents the fact that under (C3), i.e., *m* = 0, the nodes *J*_1,*t*−*m*+1_ and *J*_*m*−1,*t*−1_ vanish (adapted from Schaarschmidt (2018))

## 4 Discussion

Age- and stage-structure of fish populations may vary with time. Hence, there is a need to consider all stanzas in the life history to realistically define the transition from spawning to recruitment. Consistent with this consideration, we have presented a DTMM describing a generic life cycle of fish with four stages and several age classes. Based on this model, we have demonstrated two important principles concerning a functional presentation of the parent-progeny relationship for fish. Firstly, we have proven that in general, both the multi-variate and closed-form SR functions are limiting, because they do not track the age- and stage-structure from spawning to recruitment. Secondly, we defined necessary and sufficient conditions for the existence of a closed-form SR function.

The closest modeling approach to that presented in this paper is by Touzeau and Gouzé (1998) and Schaarschmidt et al (2018). Our work presents an extension of these previous approaches in considering all life history stages, instead of a broad classification of the population into pre-recruits (eggs, larvae, and juveniles) and adults. We have represented both the fast processes (spawning, egg and larval survival) and the slower processes (ageing of juveniles) which occur prior to recruitment.

Assuming that conditions (C2) and (C4) are true, the existence of a SR function is guaranteed by Theorem 4. These two conditions are implied when one assumes that (i) the number of individuals in the first life stage is a function of *S*, and (ii) the number of individuals in all subsequent stages is a function of the number of individuals in the previous stage. The approach by Paulik (1973) and Brooks and Powers (2007) in investigating the functional forms of parent-progeny relationships admitted by multi-stage models, was underpinned by these assumptions. It is known that cannibalism can be a limiting factor for survival (Bogstad et al 2016). Hence, inclusion of cannibalism and age-structure in our model, extends the approaches by Paulik (1973) and Brooks and Powers (2007).

We found the existence of a multi-variate SR function to be premised on either the population being at a stable fixed point, recruitment being defined as the transition rate to age class 0, or mortality of juveniles being constant with respect to the number of adults. These conditions are considered sufficient.

We proved that the existence of a multi-variate SR function is necessary, but not sufficient, for the existence of a SR function of a single variable. To ensure that the parent-progeny relationship has the conventional form of a SR function, additional conditions must be imposed on the structure of the functions representing egg production and survival of eggs and larvae. It is required that the three processes are given by functions of the same weighted sum of numbers of adults (e.g., the total number of adults or the spawning stock biomass).

When defining recruitment as the transition rate to age class 0, the evolution of all prerecruit (eggs, larvae, juveniles) happens at faster rates than aging and mortality of adult fish. This is in line with the observation made by Schaarschmidt et al (2018), who found fast prerecruit dynamics to be a prerequisite for the existence of a multi-variate SR function.

Previously, SR functions have been derived assuming age-independence of parameters related to egg production and survival of prerecruits (Quinn and Deriso 1999; Touzeau and Gouzé 1998). In this case, all age classes in the adult population contribute homogeneously to egg production and survival and the sufficient condition (C4) for the existence of an SR function holds. However, assuming age-independence of parameters is but one way of ensuring that (C4) holds.

In summary, we observe that by choosing a generic modeling framework, we found necessary and sufficient conditions for the existence and structure of the SR function with broad applicability. In this sense, our results extend and unify previous observations made under a variety of modeling assumptions.

Our results rely on a number of assumptions, e.g. discrete time transitions and unified time steps (T1 and T2). While these assumptions provide a structured foundation for our analysis, we acknowledge that the complexities of real-world fisheries populations may deviate from these simplifications.

We have considered an age-structured population though the framework can apply to populations structured by length or weight. The existence of a multi-variate SR function remains valid regardless of whether the adult population is structured according to age, length or weight. However, a multi-variate SR function in a closed-form is dependent on additional conditions.

Our results are relevant to the effective management of fisheries. Firstly, since most fish stocks exhibit complex life histories, our findings imply that they may not exhibit functional relationships that align with traditional SR functions. Hence, there is a need for a reevaluation of the applicability of traditional, closed-form SR functions in management decisions.

## 5 Data Availability

This is not applicable to this article as no datasets were generated or analyzed during the current study.

